# Retrospective Ensemble Docking of Allosteric Modulators in an Adenosine G-Protein-Coupled Receptor

**DOI:** 10.1101/2020.01.11.902809

**Authors:** Apurba Bhattarai, Jinan Wang, Yinglong Miao

## Abstract

**Background:** Ensemble docking has proven useful in drug discovery and development. It increases the hit rate by incorporating receptor flexibility into molecular docking as demonstrated on important drug targets including G-protein-coupled receptors (GPCRs). Adenosine A_1_ receptor (A_1_AR) is a key GPCR that has been targeted for treating cardiac ischemia-reperfusion injuries, neuropathic pain and renal diseases. Development of allosteric modulators, compounds binding to distinct and less conserved GPCR target sites compared with agonists and antagonists, has attracted increasing interest for designing selective drugs of the A_1_AR. Despite significant advances, more effective approaches are needed to discover potent and selective allosteric modulators of the A_1_AR.

**Methods:** Ensemble docking that integrates Gaussian accelerated molecular dynamic (GaMD) simulations and molecular docking using *Autodock* has been implemented for retrospective docking of known positive allosteric modulators (PAMs) in the A_1_AR.

**Results:** Ensemble docking outperforms docking of the receptor cryo-EM structure. The calculated docking enrichment factors (EFs) and the area under the receiver operating characteristic curves (AUC) are significantly increased.

**Conclusions:** Receptor ensembles generated from GaMD simulations are able to increase the success rate of discovering PAMs of A_1_AR. It is important to account for receptor flexibility through GaMD simulations and flexible docking.

**General Significance:** Ensemble docking is a promising approach for drug discovery targeting flexible receptors.

## Introduction

G-protein-coupled receptors (GPCRs) are the largest family of human membrane proteins. They mediate cellular responses to hormones, neurotransmitters, chemokines and the senses of sight, olfaction and taste. GPCRs have served as the primary targets for about one third of currently marketed drugs [1–3]. Particularly, the adenosine A_1_ receptor (A_1_AR) has emerged as an important therapeutic target for treating cardiac ischemia-reperfusion injuries, neuropathic pain and renal diseases[4]. However, development of the A_1_AR agonists as effective drugs has been greatly hindered. Several candidates could not progress into the clinic due to low efficacy and/or safety issues related to off-target effects. Four receptor subtypes, the A_1_, A_2A_, A_2B_ and A_3_, share a highly conserved endogenous agonist binding (“orthostatic”) site. It is challenging to design effective agonists with high selectivity.

It is appealing to design positive allosteric modulators (PAMs) that bind a less conserved, topographically distinct site and increase the responsiveness of the A_1_AR to endogenous adenosine in local regions of its production. The first selective PAM of the A_1_AR, PD81,723, was introduced in 1990[5, 6]. Since then, many research groups have performed extensive structure-activity relationship studies [6–15]. Several refined compounds were identified. Notably, T62 evaluated by King Pharmaceuticals progressed to Phase IIB clinical trial but failed due to lack of potency [16, 17]. Overall, these compounds still suffer from major limitations such as low solubility and potency for pharmaceutical use. It remains difficult to discover PAMs of higher potency for the A_1_AR.

Virtual screening has become increasingly important in the development of therapeutic drugs and discovery of novel GPCR ligands, including the antagonists, agonists, and allosteric modulators [18, 19]. Molecular docking is a widely used virtual screening technique. Early docking studies were performed using static crystal structures of target receptors and flexible ligands, which were successful in certain cases [20]. However, proteins have been often considered rigid, largely limiting the successful rate of docking. In fact, proteins are usually flexible and involve conformational changes upon ligand binding [21]. Therefore, computational approaches are needed to deal with the flexibility of proteins.

Ensemble docking of small probe molecules for flexible pharmacophore modeling was first introduced in 1999 [21]. Since then, ensemble-based [22] methods has been implemented to identify novel ligands of different proteins [23], including the immunophilin FKBP [24], HIV-1 integrase [25], RNA-editing ligase [26], membrane fusion protein [27], prothrombinase enzyme [28] and fibroblast growth factor 23 [29]. Additionally, ensemble docking has been applied to applied interactions between the drug and off-target proteins [30] and identify protein targets of natural products [31, 32].

Furthermore, ensemble docking has been applied to discover novel allosteric modulators of GPCRs. Huang et al.[33] discovered a unique PAM ogerin of GPR68 and allosteric modulators of GPR65 by docking to receptor ensembles generated from homology modeling. Long-timescale accelerated molecular dynamics (aMD) simulations were incorporated into ensemble docking to design allosteric modulators of the M2 muscarinic GPCR [34]. A number of 12 compounds with affinities ≤ 30 µM was identified, four of which were confirmed as new negative allosteric modulators (NAMs) and one as a PAM of the M2 receptor.

For the A_1_AR, X-ray structures have been determined in the inactive antagonist-bound form [35] and a cryo-EM structure in the active agonist-Gi-bound complex [36]. However, there is still no published structure of the A_1_AR bound by allosteric modulators. This has greatly hindered structure-based design of potent and selective PAMs of the A_1_AR [37]. Nevertheless, mutagenesis and molecular modeling studies have suggested that the A_1_AR allosteric site may reside within the second extracellular loop (ECL2) [38, 39]. We previously applied Gaussian accelerated molecular dynamics (GaMD) simulations to determine binding modes of two prototypical A_1_AR PAMs, PD81723 and VCP171[40]. The GaMD simulations allowed us to identify low-energy binding modes of the PAMs at an allosteric site formed by the receptor ECL2, being highly consistence with the experimental data [38, 39]. Additionally, we performed GaMD simulations on both the inactive antagonist-bound and active agonist-Gi-bound A_1_AR [41, 42]. The GPCR-membrane interactions were found to highly depend on the receptor activation state and motions in the GPCR and membrane lipids are strongly coupled [41]. These studies provided an excellent starting point for ensemble docking and rational drug design of the A_1_AR. In this study, structural ensembles were generated from GaMD simulations of the A_1_AR and used for retrospective docking of allosteric modulators (Fig. 1 and 2A). Receptor snapshots were taken every 0.2 ps from the GaMD simulations and clustered for the ECL2 target site. Ten representative structural clusters were obtained for molecular docking. We performed retrospective docking of 25 known PAMs of the A_1_AR [15, 43–46] and 2475 drug-like decoys generated from the Directory of Useful Decoys, Enhanced (DUD-E) [47]. *Autodock* was applied for both rigid-body and flexible docking [48–50]. Enrichment factors (EFs) [51] and area under the receiver operating characteristic curve (AUC) were calculated to evaluate the docking performances. Retrospective docking of the PAMs allowed us to validate structural ensembles of the A_1_AR and optimize our molecular docking protocol for future virtual screening.

**Fig. 1.**
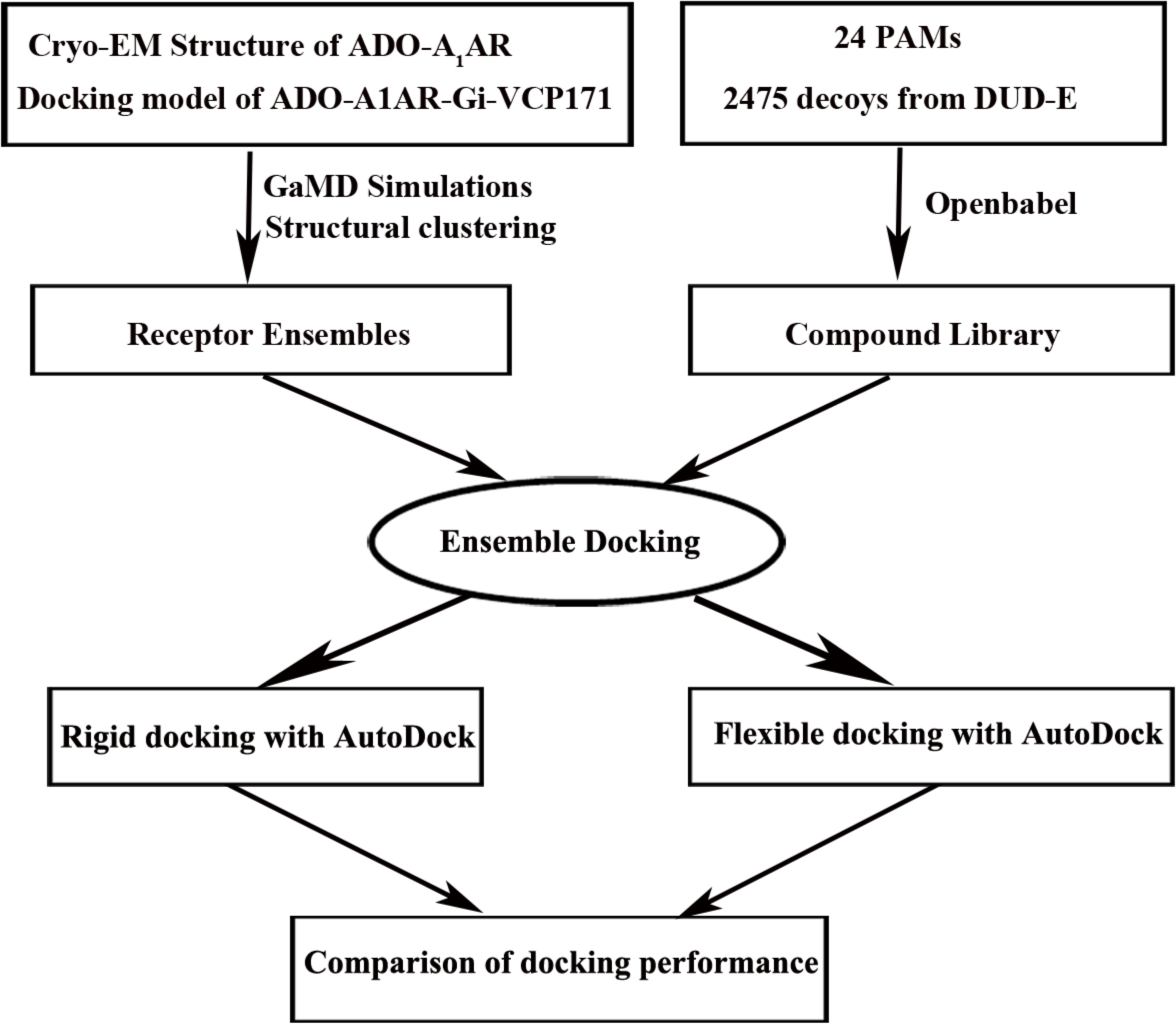
Overview flowchart for retrospective docking of positive allosteric modulators (PAMs) in the A_1_AR. Starting from the cryo-EM structure of the active ADO-Gi-bound A_1_AR (6D9H) and docking model of PAM VCP171-bound A_1_AR (ADO-A_1_AR-Gi-VCP171), GaMD simulations were carried out to construct structural ensembles to account for the receptor flexibility. Meanwhile, a compound library was prepared for 25 known PAMs of the A_1_AR and 2450 decoys obtained from the DUD-E with *openbabel* 2.4.1. Ensemble docking was then performed to identify the PAMs, for which the AUC and enrichment factors were calculated to evaluate docking performance. Both rigid-body and flexible docking were tested using *Autodock*.

**Fig. 2.**
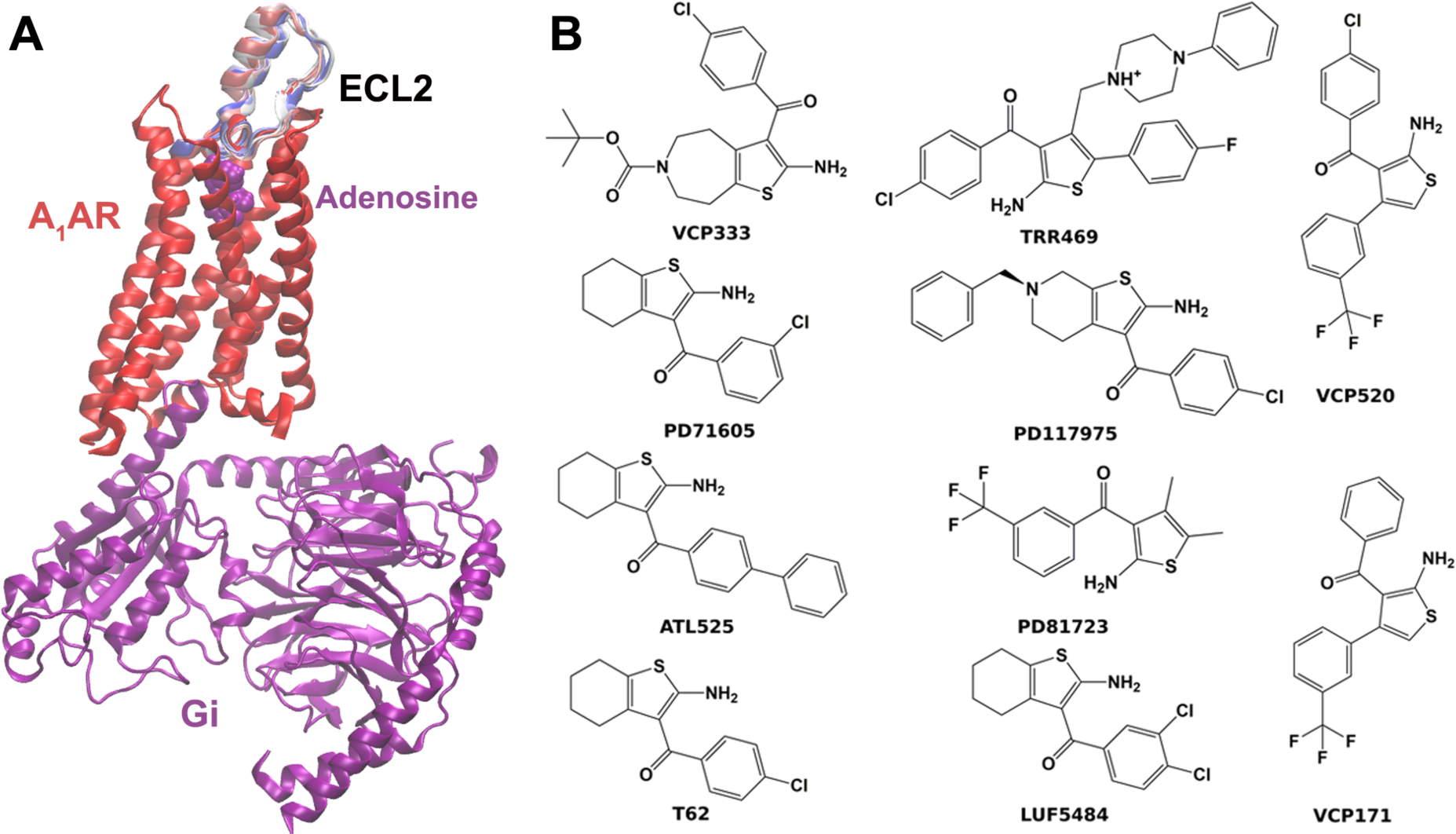
(A) Structural ensembles of the target allosteric site located in extracellular loop 2 (ECL2) generated for the A_1_AR. (B) Example PAMs of the A_1_AR used for retrospective docking of the receptor ensembles.

## Materials and Methods

### Gaussian accelerated Molecular Dynamics (GaMD)

GaMD is an enhanced sampling technique, in which a harmonic boost potential is added to reduce the system energy barriers[52]. GaMD is able to accelerate biomolecular simulations by orders of magnitude[53, 54]. GaMD does not need predefined collective variables. Moreover, because GaMD boost potential follows a Gaussian distribution, biomolecular free energy profiles can be properly recovered through cumulant expansion to the second order[52]. GaMD has successfully overcome the energetic reweighting problem in free energy calculations that was encountered in the previous accelerated molecular dynamics (aMD) method[55, 56] for free energy calculations. GaMD has been implemented in widely used software packages including AMBER [52, 57] and NAMD[58]. A brief summary of GaMD is provided here.

Consider a system with *N* atoms at positions 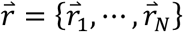. When the system potential 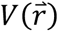 is lower than a reference energy *E*, the modified potential 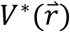 of the system is calculated as:

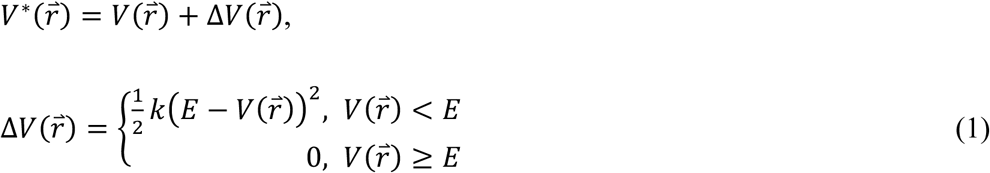

where *k* is the harmonic force constant. The two adjustable parameters *E* and *k* are automatically determined based on three enhanced sampling principles[52]. The reference energy needs to be set in the following range:

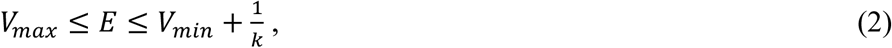

where *V*_*max*_ and *V*_*min*_ are the system minimum and maximum potential energies. To ensure that Eqn. (2) is valid, *k* has to satisfy: 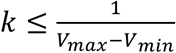 Let us define 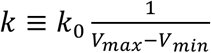, then 0 < *k*_0_ ≤ 1. The standard deviation of ∆*V* needs to be small enough (i.e., narrow distribution) to ensure proper energetic reweighting[59]: *σ*_∆*V*_ = *k*(*E* − *V*_*avg*_)*σ*_*V*_ ≤ *σ*_*0*_ where *V*_*avg*_ and *σ*_V_ are the average and standard deviation of the system potential energies, *σ*_∆*V*_ is the standard deviation of ∆*V* with *σ*_0_ as a user-specified upper limit (e.g., 10*k*_*B*_T) for proper reweighting. When *E* is set to the lower bound *E*=*V*_*max*_, *k*_0_ can be calculated as:

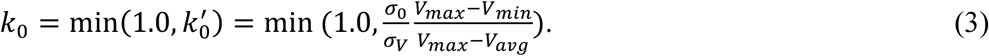

Alternatively, when the threshold energy *E* is set to its upper bound 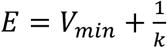 is set to:

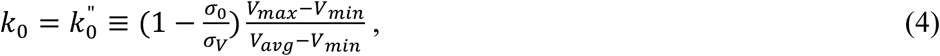

if 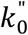 is found to be between *0* and *1*. Otherwise, *k*_0_ is calculated using Eqn. (3).

Similar to aMD, GaMD provides schemes to add only the total potential boost ∆*V*_*P*_, only dihedral potential boost ∆*V*_*D*_, or the dual potential boost (both ∆*V*_*P*_ and ∆*V*_*D*_). The dual-boost simulation generally provides higher acceleration than the other two types of simulations[60]. The simulation parameters comprise of the threshold energy E for applying boost potential and the effective harmonic force constants, *k*_0*P*_ and *k*_0*D*_ for the total and dihedral potential boost, respectively.

### Energetic reweighting of GaMD simulations

To calculate potential of mean force (PMF)[61] from GaMD simulations, the probability distribution along a reaction coordinate is written as *p*^∗^(*A*). Given the boost potential 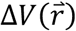 of each frame, *p*^∗^(*A*) can be reweighted to recover the canonical ensemble distribution, *p*(*A*), as:

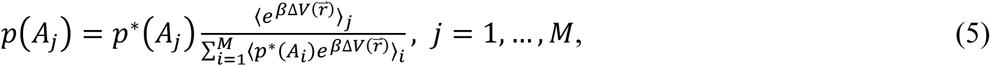

where *M* is the number of bins, 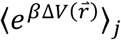 is the ensemble-averaged Boltzmann factor of 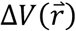 for simulation frames found in the *j*^th^ bin. The ensemble-averaged reweighting factor can be approximated using cumulant expansion:

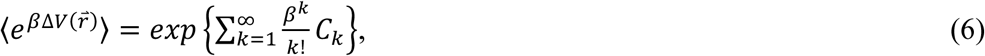

where the first two cumulants are given by

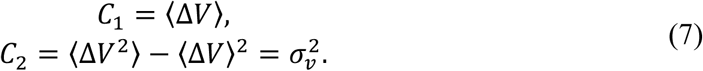

The boost potential obtained from GaMD simulations usually follows near-Gaussian distribution. Cumulant expansion to the second order thus provides a good approximation for computing the reweighting factor[52, 59]. The reweighted free energy *F*(*A*) = −*k*_*B*_*T* ln *p*(*A*) is calculated as:

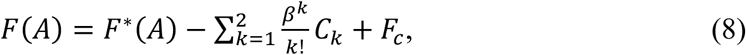

where *F*^∗^(*A*) = −*k*_*B*_*T* ln *p*^∗^(*A*) is the modified free energy obtained from GaMD simulation and *F*_*c*_ is a constant.

### System setup

The cryo-EM structure of the ADO-A_1_AR-Gi complex (PDB: 6D9H[36]) were used to prepare the simulation systems. In order to prepare PAM-bound structures of the active A_1_AR for docking, the VCP171 was docked with *Autodock* to the ECL2 allosteric site of the A_1_AR [38–40]. Then the binding conformation of VCP171 with the highest docking score was chosen as initial structure of the PAM-bound A_1_AR. For both PAM-free (*apo*) and PAM-bound (*holo*) structures of the A_1_AR, all chain termini were capped with neutral groups, i.e. the acetyl group (ACE) for the N-terminus and methyl amide group (CT3) for C terminus. Protein residues were set to the standard CHARMM protonation states at neutral pH with the *psfgen* plugin in VMD[62]. Then the receptor was inserted into a POPC bilayer with all overlapping lipid molecules removed using the *Membrane* plugin in VMD[62]. The system charges were then neutralized at 0.15 M NaCl using the *Solvate* plugin in VMD[62]. Periodic boundary conditions were applied on the simulation systems.

The CHARMM36 parameter set [63] was used for the protein and POPC lipids. For agonist ADO and PAM VCP171, the force field parameters were obtained from the CHARMM ParamChem web server[38, 64]. Initial energy minimization and thermalization of the A_1_AR system follow the same protocol as used in the previous GPCR simulations[65]. The simulation proceeded with equilibration of lipid tails. With all the other atom fixed, the lipid tails were energy minimized for 1000 steps using the conjugate gradient algorithm and melted with constant number, volume, and temperature (NVT) run for 0.5ns at 310 K. Each system was further equilibrated using constant number, pressure, and temperature (NPT) run at 1 atm and 310 K for 10 ns with 5 kcal (mol Å^2^)^−1^ harmonic position restraints applied to the protein. Further equilibration of the systems was performed using an NPT run at 1 atm and 310 K for 0.5ns with all atoms unrestrained. Conventional MD simulation was performed on each system for 10 ns at 1atm pressure and 310 K with a constant ratio constraint applied on the lipid bilayer in the X-Y plane. Both dihedral and dual-boost GaMD simulations (GaMD_Dih and GaMD_dual respectively) were then performed using NAMD2.13[58, 66] and AMBER 18 [52] to generate receptor ensembles for docking. Simulations of the A_1_AR were summarized in Table 1. In the GaMD simulations, the threshold energy *E* for adding boost potential is set to the lower bound, i.e. *E = V*_*max*_[52, 58]. The simulations included 50ns equilibration after adding the boost potential and then three independent production runs lasting 300 *ns* with randomized initial atomic velocities. GaMD production simulation frames were saved every 0.2ps. Snapshots of all three GaMD production simulations were combined for clustering to identify representative structures for docking using the hierarchical agglomerative algorithm in CPPTRAJ[67]. The *PyReweighting* toolkit[59] was applied to reweight GaMD simulations by combining independent trajectories for each system. Free energy values were calculated for the top-ranked structural clusters of the receptor.

**Table 1:** Summary of GaMD simulations performed on the adenosine A_1_ receptor (A_1_AR).

| System | <sup>a</sup>$N_{\text{atoms}}$ | Dimension($\text{\AA}^3$) | Program | <sup>b</sup>Method | Simulation | <sup>c</sup>$\Delta V_{\text{avg}}$<br>(kcal/mol) | <sup>d</sup>$\sigma_{\Delta V}$<br>(kcal/mol) |
| --- | --- | --- | --- | --- | --- | --- | --- |
| ADO-A<sub>1</sub>AR-Gi | 180,394 | 93x111x167 | NAMD | GaMD_Dih | 300 ns x 2 | 5.04 | 2.22 |
|  |  |  |  | GaMD_Dual | 300 ns x 2 | 10.14 | 2.59 |
| ADO-A<sub>1</sub>AR-Gi | 180,394 | 93x111x167 | AMBER | GaMD_Dih | 300 ns x 3 | 6.89 | 4.06 |
|  |  |  |  | GaMD_Dual | 300 ns x 3 | 21.43 | 6.50 |
| ADO-A<sub>1</sub>AR-Gi-VCP171 | 180,403 | 93x111x167 | AMBER | GaMD_Dih | 300 ns x 3 | 6.80 | 2.74 |
|  |  |  |  | GaMD_Dual | 300 ns x 3 | 24.80 | 6.61 |

<sup>a</sup> $N_{\text{atoms}}$ is number of atoms in the simulation systems.
<sup>b</sup>GaMD\_Dih and GaMD\_Dual represent the dihedral and dual boost GaMD simulations respectively.
<sup>c</sup> $\Delta V_{\text{avg}}$ and <sup>d</sup> $\sigma_{\Delta V}$ are the average and standard deviation of the GaMD boost potential.

### Retrospective docking of known allosteric ligands to the A_1_AR

A number of 25 known PAMs of the A_1_AR were collected from literature [15, 43–46] (Fig. 2B) and 2475 decoys were obtained from the DUD-E database [47]. This compound library served as a dataset to validate the receptor ensembles constructed from GaMD simulations of the A_1_AR. Docking was performed on these compounds to the receptor ECL2 site using *Autodock*[48]. Default parameters (number of individuals in population (ga_pop_size), 1,500; maximum number of generations (ga_num_generations), 27,000 and number of genetic algorithms run (ga_run), 10) were used for the *Autodock* docking calculations unless described otherwise. A set of docking algorithms were extensively tested, including the flexible docking and rigid-body docking at different levels (the long level with ga_num_evals of 25,000,000, medium level with ga_num_evals of 2,500,000 and short level with ga_num_evals of 250,000). For the flexible docking, residues within 5 Å of VCP171 in the docking pose were selected as flexible residues. The representative structures obtained from GaMD simulations and the cryo-EM structure of the A_1_AR (PDB ID: 6D9H) were used for the receptor. The ADO agonist was kept in all the docking calculations. Firstly, all polar hydrogens were added and Gasteiger charges were assigned to atoms in the receptor. Secondly, a 3D search grid was created with the *AutoGrid* algorithm[68] to calculate binding energies of the ligands and decoys in the A_1_AR. The center of mass of the receptor ECL2 was chosen as the grid center and a box size of 60*60*60 Å^3^ was applied for docking. The Lamarckian genetic algorithm[68] was applied to model protein-ligand interactions.

For docking of GaMD simulation structural clusters, the predicted binding energy for each conformation was calculated by its raw docking score and reweighted according to the following:

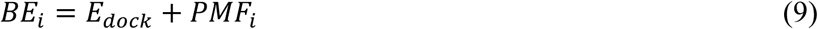

where *PMF*_*i*_ is the reweighted free energy of the *i*th receptor structural cluster and *E*_*dock*_ is the corresponding raw docking score. Ranking of the docked compounds was examined in terms of both the minimum predicted binding energy obtained for each compound against any of the given receptor structures (“*BE*_*min*_”) and the average of predicted binding energies (“*BE*_*avg*_”) [34]. The average binding energy is calculated as:

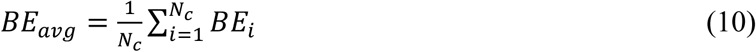

where *Nc* is the total number of receptor structural clusters, *BE*_*i*_ is the binding energy of a compound to the *i*th cluster. The minimum predicted binding energy is given by:

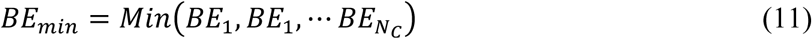

Enrichment factor (EF) is calculated by:

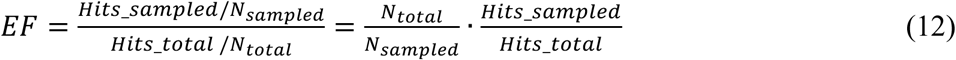

where the *N*_*total*_, *N*_*sampled*_, *Hits_total* and *Hits_sampled* are the number of total compounds in the database, number of compounds for ranking, number of total hits in the database and number of hits found among the ranked compounds, respectively. The “early” enrichment factor (EF’) that accounts for the rank of each of the *Hits_sampled* known actives is calculated by:

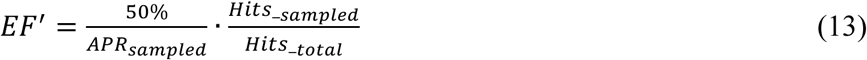

Where the *APR*_*sampled*_ is the average percentile rank of the *Hits_sampled* known actives.

## Results

### Construction of receptor ensembles through structural clustering of GaMD simulations

All-atom GaMD simulations were performed on different systems of the active A_1_AR with varied levels of acceleration, i.e. the dihedral and dual boost (see **Methods** and Table 1). Two different software packages, NAMD2.13[58, 66] and AMBER 18[52, 57] were used for the simulations. Six sets of GaMD simulations, including the ADO-A_1_AR-Gi (NAMD, GaMD_Dih), ADO-A_1_AR-Gi (NAMD, GaMD_Dual), ADO-A_1_AR-Gi (AMBER, GaMD_Dih), ADO-A_1_AR-Gi (AMBER, GaMD_Dual), ADO-A_1_AR-Gi-VCP171 (AMBER, GaMD_Dih) and ADO-A_1_AR-Gi-VCP171 (AMBER, GaMD_Dual) were obtained to generate receptor ensembles. Overall, GaMD_Dual provided higher boost potential than the GaMD_Dih. AMBER provided higher boost potential than NAMD in the GaMD simulations due to different algorithms used to calculate the potential average and standard deviation[40]. The GaMD_Dual simulations of ADO-A_1_AR-Gi with AMBER recorded an average boost potential of 21.43 kcal/mol and 6.50 kcal/mol standard deviation compared with 10.14 kcal/mol average and 2.59 kcal/mol standard deviation in the GaMD_Dual simulations with NAMD. The GaMD_Dih simulations of ADO-A_1_AR-Gi using AMBER also exhibited higher average boost potential of 6.89 kcal/mol and 4.06 kcal/mol standard deviation compared with 5.04 kcal/mol average and 2.22 kcal/mol standard deviation in the GaMD_Dih simulations using NAMD. For the VCP171 bound A_1_AR system, the GaMD_Dual simulations with AMBER recorded an 24.80 kcal/mol average boost potential and 6.61 kcal/mol standard deviation compared with the average of 6.80 kcal/mol and of and 2.47 kcal/mol standard deviation in the GaMD_Dih simulations with AMBER.

Ten representative structures of the receptor ECL2 allosteric site were obtained by root-mean-square deviation (RMSD)-based structural clustering of the receptor snapshots for each set of the GaMD simulations. The representative structural clusters were characterized by the fraction of simulation frames in the cluster, the GaMD reweighted PMF free energy values of each cluster, and their sampled conformational space (Table 2). Overall, the top clusters in each receptor ensemble contributed to significantly higher fractions of simulation frames than the lower ranked clusters. However, the free energy values were ordered differently for most of the structural clusters with GaMD reweighting. It was thus important to take GaMD reweighted free energies into account in the ensemble docking. Notably, structural clusters obtained from AMBER GaMD_Dual simulations of ADO-A_1_AR-Gi-VCP171 system sampled the largest conformational space, as characterized by ~4.5-5.9 Å average distance between the cluster centroids (AvgCDist) and 2.31 Å average distance between simulation frames (AvgDist) of the lowest free energy/top-ranked cluster (Table 2). In comparison, clusters from AMBER GaMD_Dih simulations of the VCP171-bound A_1_AR sampled the smallest conformational space with ~2.1-2.4 Å average cluster centroid distance and 1.78 Å average distance between simulation fames in the top cluster. Clusters obtained from GaMD simulations of the ADO-A_1_AR-Gi system sampled conformational space in the medium range (Table 2). These receptor structural ensembles were subsequently used for retrospective docking of PAMs in the A_1_AR.

**Table 2:** Summary of receptor structural clusters obtained from different GaMD simulations of the A_1_AR. Ten representative structural clusters are listed with their fraction of simulation frames, the GaMD reweighted PMF free energy values (kcal/mol), the average distance of each cluster centroid to every other cluster centroid (AvgCDist) and the average distance between frames in the cluster (AvgDist).

| Cluster ID | Cluster Properties | ADO-A<sub>1</sub>AR-Gi (NAMD, GaMD_Dih) | ADO-A<sub>1</sub>AR-Gi (NAMD, GaMD_Dual) | ADO-A<sub>1</sub>AR-Gi (AMBER, GaMD_Dih) | ADO-A<sub>1</sub>AR-Gi (AMBER, GaMD_Dual) | ADO-A<sub>1</sub>AR-Gi-VCP171 (AMBER, GaMD_Dih) | ADO-A<sub>1</sub>AR-Gi-VCP171 (AMBER, GaMD_Dual) |
| --- | --- | --- | --- | --- | --- | --- | --- |
| Cluster 1 | Fraction | 0.22 | 0.18 | 0.50 | 0.78 | 0.73 | 0.56 |
|  | Reweighted PMF (kcal/mol) | 0.19 | 1.78 | 0.00 | 1.64 | 0.00 | 0.00 |
|  | AvgCDist (Å) | 3.01 | 3.50 | 4.18 | 2.76 | 2.10 | 4.62 |
|  | AvgDist (Å) | 2.40 | 2.29 | 2.28 | 1.82 | 1.78 | 2.31 |
| Cluster 2 | Fraction | 0.16 | 0.16 | 0.16 | 0.09 | 0.15 | 0.16 |
|  | Reweighted PMF (kcal/mol) | 0.40 | 1.27 | 0.80 | 2.83 | 0.97 | 1.40 |
|  | AvgCDist (Å) | 3.26 | 3.04 | 4.51 | 3.02 | 2.37 | 4.51 |
|  | AvgDist (Å) | 2.30 | 2.40 | 1.99 | 1.69 | 1.64 | 2.62 |
| Cluster 3 | Fraction | 0.15 | 0.14 | 0.14 | 0.04 | 0.07 | 0.14 |
|  | Reweighted PMF (kcal/mol) | 1.11 | 0.00 | 0.61 | 3.52 | 1.45 | 0.35 |
|  | AvgCDist (Å) | 3.05 | 3.12 | 3.75 | 2.97 | 2.24 | 5.41 |
|  | AvgDist (Å) | 2.35 | 2.31 | 2.60 | 1.88 | 1.62 | 2.73 |
| Cluster 4 | Fraction | 0.14 | 0.13 | 0.11 | 0.04 | 0.02 | 0.05 |
|  | Reweighted PMF (kcal/mol) | 1.14 | 1.41 | 1.19 | 3.70 | 1.60 | 2.93 |
|  | AvgCDist (Å) | 3.03 | 3.57 | 3.94 | 3.42 | 2.33 | 4.80 |
|  | AvgDist (Å) | 2.38 | 2.20 | 2.25 | 1.74 | 1.57 | 2.65 |
| Cluster 5 | Fraction | 0.10 | 0.12 | 0.04 | 0.03 | 0.02 | 0.03 |
|  | Reweighted PMF (kcal/mol) | 1.38 | 1.25 | 1.99 | 3.41 | 1.83 | 2.09 |
|  | AvgCDist (Å) | 2.98 | 3.10 | 3.77 | 3.11 | 2.34 | 4.60 |
|  | AvgDist (Å) | 2.23 | 2.25 | 2.00 | 1.66 | 1.35 | 2.78 |
| Cluster 6 | Fraction | 0.07 | 0.06 | 0.03 | 0.01 | 0.004 | 0.02 |
|  | Reweighted PMF (kcal/mol) | 2.75 | 0.85 | 1.54 | 4.15 | 2.93 | 2.79 |
|  | AvgCDist (Å) | 2.95 | 3.31 | 4.62 | 3.06 | 2.22 | 4.76 |
|  | AvgDist (Å) | 2.14 | 2.19 | 2.47 | 1.69 | 1.49 | 2.30 |
| Cluster 7 | Fraction | 0.06 | 0.06 | 0.01 | 0.001 | 0.004 | 0.02 |
|  | Reweighted PMF (kcal/mol) | 2.23 | 0.71 | 1.82 | 4.86 | 3.41 | 1.17 |
|  | AvgCDist (Å) | 3.12 | 3.51 | 4.34 | 3.07 | 2.22 | 5.42 |
|  | AvgDist (Å) | 2.33 | 2.35 | 2.23 | 1.74 | 1.71 | 2.45 |
| Cluster 8 | Fraction | 0.06 | 0.06 | 0.004 | 0.00 | 0.00 | 0.01 |
|  | Reweighted PMF (kcal/mol) | 2.18 | 2.32 | 2.38 | 5.69 | 3.36 | 1.86 |
|  | AvgCDist (Å) | 3.13 | 3.40 | 4.01 | 2.73 | 2.22 | 5.33 |
|  | AvgDist (Å) | 2.28 | 2.16 | 1.63 | 1.41 | 1.42 | 1.88 |
| Cluster 9 | Fraction | 0.03 | 0.05 | 0.003 | 0.001 | 0.001 | 0.01 |
|  | Reweighted PMF (kcal/mol) | 2.28 | 1.02 | 3.43 | 0.73 | 4.01 | 4.10 |
|  | AvgCDist (Å) | 3.03 | 3.38 | 3.83 | 3.36 | 2.24 | 4.78 |
|  | AvgDist (Å) | 2.11 | 2.10 | 1.60 | 0.00 | 1.19 | 1.73 |
| Cluster 10 | Fraction | 0.02 | 0.03 | 0.002 | 0.001 | 0.001 | 0.00 |
|  | Reweighted PMF (kcal/mol) | 0.00 | 1.07 | 4.77 | 0.00 | 2.86 | 5.78 |
|  | AvgCDist (Å) | 3.90 | 3.09 | 3.77 | 3.57 | 2.22 | 5.87 |
|  | AvgDist (Å) | 2.22 | 2.14 | 1.65 | 0.00 | 1.35 | 0.00 |

### Correction with GaMD reweighting improved retrospective docking performances

The GaMD receptor ensembles were used for docking of the compound library consisting of 25 known PAMs and 2475 decoys of the A_1_AR. The EF and EF’ enrichment factors and AUC values were calculated to evaluate the ensemble docking performances [51]. The EF and EF’ were calculated for top 5% and 10% of total sampled compounds (Table 3). For the ten representative conformations obtained from each set of GaMD simulations, both the minimum binding energy (*BE*_*min*_) and the average binding energy (*BE*_*avg*_) were used to assess the docking performances. To evaluate the effects of GaMD reweighting, we compared the binding energies using both the raw scores obtained from docking calculations (“**raw”** in Table 3) and the reweighted scores corrected with cluster free energy values calculated by GaMD reweighting **(“reweighted”** in Table 3). For a total of 60 calculated docking metrics regarding the AUC, EF (5%), EF (10%), EF’ (5%) and EF’ (10%) as listed in Table 3, 32 (53.3%) of them showed better performance using the GaMD-reweighted scores than using the raw scores. While only 10 (16.7%) of them showed decreased performance and 18 (30%) with the same performance (Table 3).

**Table 3:** Summary of docking results for the AUC and the EF and EF’ values with 5% and 10% of sampled compounds calculated using different GaMD simulation ensembles of the A_1_AR. Among the listed 60 docking performances metrics, 32 of them (highlighted in bold) showed better performance using the GaMD-reweighted scores than using the raw scores, while only 18 and 10 of them showed the same and decreased performance values, respectively.

| System | Ranking | AUC |  | EF (5%) |  | EF' (5%) |  | EF (10%) |  | EF' (10%) |  |
| --- | --- | --- | --- | --- | --- | --- | --- | --- | --- | --- | --- |
|  |  | raw | reweighted | raw | reweighted | raw | raw | reweighted | raw | reweighted | raw |
| ADO-A<sub>1</sub>AR-Gi (NAMD, GaMD_Dih) | $BE_{avg}$ | 0.48 | 0.45 | 0.00 | 0.00 | 0.51 | **0.78** | 0.00 | **0.92** | 0.00 | **0.96** |
| | $BE_{min}$ | 0.46 | **0.51** | 0.00 | 0.00 | 0.77 | 0.46 | 0.00 | **1.09** | 0.40 | **0.48** |
| ADO-A<sub>1</sub>AR-Gi (NAMD, GaMD_Dual) | $BE_{avg}$ | 0.45 | 0.45 | 0.00 | 0.00 | 0.78 | 0.78 | 0.00 | **0.96** | 0.78 | **0.96** |
| | $BE_{min}$ | 0.46 | **0.51** | 0.00 | 0.00 | 1.47 | 0.46 | 0.80 | **2.47** | 0.46 | **0.48** |
| ADO-A<sub>1</sub>AR-Gi (AMBER, GaMD_Dih) | $BE_{avg}$ | 0.44 | **0.61** | 0.00 | 0.00 | 0.36 | **3.73** | 0.00 | **0.71** | 1.20 | **5.03** |
| | $BE_{min}$ | 0.43 | **0.49** | 0.00 | 0.00 | 0.42 | **0.53** | 0.00 | **0.76** | 0.00 | **0.97** |
| ADO-A<sub>1</sub>AR-Gi (AMBER, GaMD_Dih) | $BE_{avg}$ | 0.43 | 0.43 | 0.00 | 0.00 | 0.48 | 0.47 | 0.00 | **0.71** | 0.00 | **0.71** |
| | $BE_{min}$ | 0.42 | 0.35 | 0.00 | 0.00 | 0.91 | 0.42 | 0.00 | **1.10** | 0.00 | **0.67** |
| ADO-A<sub>1</sub>AR-Gi-VCP171 (AMBER, GaMD_Dih) | $BE_{avg}$ | 0.47 | 0.47 | 0.00 | 0.00 | 0.42 | 0.42 | 0.00 | **0.77** | 0.00 | **0.77** |
| | $BE_{min}$ | 0.49 | **0.50** | 0.00 | 0.00 | 1.48 | 0.60 | 0.40 | **1.53** | 0.00 | **1.10** |
| ADO-A<sub>1</sub>AR-Gi-VCP171 (AMBER, GaMD_Dual) | $BE_{avg}$ | 0.47 | 0.44 | 0.00 | 0.00 | 0.37 | 0.37 | 0.00 | **0.67** | 0.00 | **0.66** |
| | $BE_{min}$ | 0.49 | 0.46 | 0.00 | 0.00 | 0.73 | 0.38 | 0.00 | **1.10** | 0.00 | **0.73** |

With GaMD reweighting, the calculated AUC using ranking by the minimum binding energy (*BE*_*min*_) increased by ~10-14% for receptor ensembles obtained from NAMD GaMD_Dih, NAMD GaMD_Dual and AMBER GaMD_Dih simulations of the ADO-A_1_AR-Gi system. The AUC using ranking by the average binding energy (*BE*_*avg*_) increased from 0.44 to 0.61 for receptor ensemble from AMBER GaMD_Dih of the ADO-A_1_AR-Gi system. More importantly, while the EF and EF’ values for 5% of sampled compounds were generally small for both raw and reweighted docking scores, the EF and EF’ for 10% of sampled compound increased significantly with reweighted scores for receptor ensembles of all the simulations (Table 3). This would be more relevant for drug design projects since only top ranked compounds could often be tested in experiments. Therefore, binding energies corrected by GaMD reweighting improved the docking performances.

Furthermore, we compared the calculated AUC, EF and EF’ in terms of ranking using the minimum binding energy (*BE*_*min*_) and average binding energies (*BE*_*avg*_). Although with exceptions, the usage of *BE*_*avg*_ mostly outperformed that of *BE*_*min*_, especially in the calculated EF’ (5%) and EF’ (10%) (Table 3). Notably, the EF’(5%) increased by ~70% to 7 times using *BE*_*avg*_ in ensemble docking of the ADO-A_1_AR-Gi system from the NAMD GaMD_Dih, NAMD GaMD_Dual and AMBER GaMD_Dih simulations than using the *BE*_*min*_, and ~2-5 times for the EF’(10%) (Table 3). The average predicted binding energy (*BE*_*avg*_) was thus applied for further parameter testing of the docking calculations.

### Flexible docking performed better than different levels of rigid-body docking

Next, we compared the performances of flexible docking and different levels of rigid-body docking with *Autodock* (Table 4). In a total of 30 calculated docking metrics regarding the AUC, EF(5%), EF(10%), EF’(5%) and EF’(10%), 22 (73.3%) of them showed improved performance with flexible docking compared with rigid-body docking at the short, medium and long levels, while only 2 (6.7%) showed the same performance and 6 (20%) with decreased performance. Among the 22 metrics that showed improved performance with flexible docking, 17 of these values increased by more than 2 times compared with rigid-body docking (Table 4).

**Table 4:** Summary of docking results for the flexible and rigid-body docking at the “short”, “medium” and “long” levels using *Autodock* regarding the AUC, EF and EF’ values with 5% and 10% of sampled compounds calculated using different GaMD simulation ensembles of the A_1_AR. Among the listed 30 docking performance metrics, 22 of them (highlighted in bold) showed better performance using flexible docking than using rigid docking, while only 2 and 6 of them showed the same and decreased performance values, respectively.

| System | Levels of Docking | ADO-A<sub>1</sub>AR-Gi (NAMD, GaMD_Dih) | ADO-A<sub>1</sub>AR-Gi (NAMD, GaMD_Dual) | ADO-A<sub>1</sub>AR-Gi (AMBER, GaMD_Dih) | ADO-A<sub>1</sub>AR-Gi (AMBER, GaMD_Dual) | ADO-A<sub>1</sub>AR-Gi-VCP171 (AMBER, GaMD_Dih) | ADO-A<sub>1</sub>AR-Gi-VCP171 (AMBER, GaMD_Dual) |
| --- | --- | --- | --- | --- | --- | --- | --- |
| AUC | Short | 0.45 | 0.45 | 0.61 | 0.43 | 0.47 | 0.44 |
|  | Medium | 0.49 | 0.49 | 0.67 | 0.50 | 0.50 | 0.48 |
|  | Long | 0.50 | 0.50 | 0.23 | 0.54 | 0.50 | 0.52 |
|  | Flexible | **0.53** | **0.53** | 0.53 | 0.51 | **0.54** | **0.60** |
| EF (5%) | Short | 0.00 | 0.00 | 0.80 | 0.00 | 0.00 | 0.00 |
|  | Medium | 0.00 | 0.00 | 3.20 | 0.00 | 0.00 | 0.00 |
|  | Long | 0.00 | 0.00 | 0.80 | 0.00 | 0.00 | 0.00 |
|  | Flexible | **0.80** | **0.80** | 0.00 | **0.80** | **0.80** | **0.80** |
| EF’ (5%) | Short | 0.78 | 0.78 | 3.73 | 0.47 | 0.42 | 0.37 |
|  | Medium | 0.95 | 0.95 | 4.81 | 0.89 | 1.11 | 1.14 |
|  | Long | 0.10 | 0.10 | 3.52 | 1.36 | 0.87 | 0.88 |
|  | Flexible | **5.00** | **5.00** | 1.41 | **6.58** | **3.97** | **8.33** |
| EF (10%) | Short | 0.00 | 0.00 | 1.20 | 0.00 | 0.00 | 0.00 |
|  | Medium | 0.00 | 0.00 | 0.40 | 0.00 | 0.40 | 0.40 |
|  | Long | 0.00 | 0.00 | 2.00 | 0.40 | 0.00 | 0.00 |
|  | Flexible | **0.40** | **0.40** | 0.40 | 0.40 | 0.40 | **0.80** |
| EF’ (10%) | Short | 0.96 | 0.96 | 5.03 | 0.71 | 0.77 | 0.66 |
|  | Medium | 1.12 | 1.12 | 7.41 | 1.06 | 1.16 | 1.13 |
|  | Long | 0.20 | 0.20 | 2.72 | 1.69 | 1.11 | 1.03 |
|  | Flexible | **3.05** | **3.05** | 1.42 | **2.46** | **2.32** | **4.98** |

The AUC increased with flexible docking of all the receptor ensembles except the ADO-A_1_AR-Gi ensemble from the AMBER GaMD_Dih and GaMD_Dual simulations (Table 4). The EF (5%) increased from 0.0 with different levels of rigid-body docking to 0.8 with flexible docking for all the receptor ensembles except the ADO-A_1_AR-Gi ensemble from the AMBER GaMD_Dih simulations. Similar significant increase was observed for the EF’ (5%) values with flexible docking of these receptor ensembles. Notably, AMBER GaMD_Dual simulations of the PAM VCP171-bound ADO-A_1_AR-Gi and PAM-free ADO-A_1_AR-Gi systems combined with flexible docking provided the highest EF’ (5%) for, i.e., 8.33 and 6.58, respectively. For the EF (10%) and EF’(10%), flexible docking led to significantly higher performance values for most of the simulations receptor ensembles than the rigid-body docking (Table 4).In summary, flexible docking provided significantly improved performances than rigid-body docking, suggesting that the flexibility of protein side chains played an important role even in the ensemble docking calculations.

### Optimized ensemble docking protocol and comparison with docking using cryo-EM structure

Comparing flexible docking of all six receptor ensembles with ranking by the GaMD reweighted the average binding energy (*BE*_*avg*_), we found that the AMBER GaMD_Dual simulations of the PAM VCP171 bound ADO-A_1_AR-Gi system provided the best docking performance (Table 4). The GaMD simulations of VCP171-bound A_1_AR sampled more relevant receptor conformations that favored binding of the PAMs, although the ECL2 allosteric site was highly flexible [41, 42] and the VCP171 PAM exhibited high fluctuations. Such conformations were not well sampled in GaMD simulations of the PAM-free (*apo*) A_1_AR started from the cryo-EM structure of ADO-A_1_AR-Gi.

For comparison, we performed flexible docking using *Autodock* with only the cryo-EM structure of the active ADO-A_1_AR-Gi (PDB: 6D9H). While the EF(5%) and EF(10%) of flexible docking of the cryo-EM structure exhibited the same values as ensemble docking of the PAM-bound A_1_AR, the AUC, EF’(5%) and EF’(10%) values decreased significantly to 0.53, 2.88 and 3.47, respectively, compared to the latter (Table 5). It is worth noting that the performance of docking the PAMs in the current study was relatively lower than that of the orthosteric ligands that bind inside class A GPCRs with generally higher affinities [34]. Nonetheless, with GaMD simulations and flexible docking, we have optimized our ensemble docking protocol to effectively account for the receptor flexibility, which will facilitate virtual screening of PAMs for the A_1_AR.

**Table 5:** Summary of docking results for the AUC and the EF and EF’ values in 5% and 10% of sampled compounds calculated using the finally selected ensemble docking model and cryo-EM structure of the A_1_AR.

| System | AUC | EF (5%) | EF' (5%) | EF (10%) | EF' (10%) |
| --- | --- | --- | --- | --- | --- |
| ADO-A<sub>1</sub>AR-Gi-VCP171 (AMBER, GaMD_Dual) | 0.60 | 0.80 | 8.33 | 0.80 | 4.98 |
| ADO-A<sub>1</sub>AR-Gi (Cryo-EM, PDB:6D9H) | 0.53 | 0.80 | 2.88 | 0.80 | 3.47 |

## Discussion

In this study, we have carried out extensive retrospective docking calculations of PAM binding to the A_1_AR using GaMD enhanced simulations and *AutoDock*. The GaMD simulations have been performed using the AMBER and NAMD simulation packages at different acceleration levels (dihedral and dual boost). The rigid-body docking at different short, medium and long levels and flexible docking have been all evaluated in our ensemble docking protocol.

Overall, flexible docking performed significantly better than the rigid-body docking at different levels with *AutoDock*. This suggested that the flexibility of protein side chains in ensemble docking is also important. The side chains of representative receptor structures obtained from GaMD simulations might be still in unfavored conformations for PAM binding. Flexible docking of protein side chains could then alleviate this problem to achieve better performance. The Glide induced fit docking was also found to outperform rigid-body docking in a previous study to identify allosteric modulators of the M2 muscarinic GPCR [34]. Generally speaking, flexible docking greatly helps accounting for protein flexibility in the side chains and improves docking performances[69].

To fully account for the protein flexibility, especially the backbone, we further incorporated GaMD enhanced sampling simulations in an ensemble docking protocol (Fig. 1). In the GaMD simulations, the boost potential obtained for one system with AMBER was greater than with NAMD due to different algorithms used to calculate the system potential statistics[40]. Accordingly, larger conformation space of the receptor was sampled in the AMBER simulations, which resulted in improved docking performance. Correction of binding energy by the GaMD reweighted free energy of the receptor structural cluster improved the docking performances. With GaMD reweighted scores, ranking by the average binding energy (*BE*_*avg*_) performed better than by the minimum binding energy (*BE*_*min*_) in terms of the AUC, EF and EF’ docking metrics.

Receptor ensemble obtained from AMBER GaMD_Dual simulations of the VCP171 PAM-bound ADO-A_1_AR-Gi outperformed other receptor ensembles for docking. This ensemble consists of snapshots of the *holo* A_1_AR with PAM bound at the ECL2 allosteric site. Interactions between the PAM and receptor ECL2 could induce more suitable conformations for PAM binding, which were otherwise difficult to sample in the simulations of PAM-free (*apo*) A_1_AR. Dual-boost GaMD was observed to perform better than the dihedral-boost GaMD for ensemble docking, suggesting that GaMD at the higher dual-boost acceleration level was needed to sufficiently sample conformational space of the GPCR PAM binding site. This finding is in contrast to the previous docking study of allosteric modulators to the M_2_ receptor that showed dihedral-boost aMD outperformed dual-boost aMD for ensemble docking [34]. Such discrepancy likely resulted from the relatively lower boost potential with a new formula applied in the GaMD simulations compared with the previous aMD method [52, 58]. The aMD simulations especially with dual boost appeared to provide too high acceleration and sample receptor conformations that did not facilitate compound docking. In comparison, dual-boost GaMD generated more appropriate receptor ensembles for docking. GaMD also solved the energetic noise problem of aMD in the protein simulations. The reweighted free energies of receptor structural clusters obtained from GaMD simulations have been shown to further improve ranking of the compounds and ensemble docking performance as demonstrated here.

In summary, we have optimized the protocol for ensemble docking of allosteric modulators to the A_1_AR, a prototypical GPCR. The ensemble docking integrates all-atom dual-boost GaMD simulations of the PAM-bound (*holo*) agonist-A_1_AR-Gi complex, flexible docking with *AutoDock* and compound scoring with the GaMD reweighted average binding energy. Enhanced sampling simulations and flexible docking have greatly improved the docking performance by effectively accounting for the protein flexibility in both the backbone and side chains. Such an ensemble docking protocol will greatly facilitate future allosteric drug design of the A_1_AR and other GPCRs.

## CRediT author statement

**Apurba Bhattarai**: Methodology, Software, Validation, Investigation, Data curation, Writing-Original Draft, Visualization. **Jinan Wang**: Methodology, Software, Validation, Investigation, Data curation, Writing-Original Draft, Visualization. **Yinglong Miao**: Conceptualization, Methodology, Software, Resources, Writing - Review & Editing, Supervision, Project administration, Funding acquisition.

## Conflict of Interest Statement

There is no conflict of interest.

## Acknowledgements

We would like to dedicate this manuscript to the 60^th^ birthday of Prof. Jeremy Smith. We appreciate the preliminary docking work of Shulammite Lim and thank Prof. Arthur Christopoulos, Dr. Lauren May, Dr. Anh Nguyen and Prof. Jens Carlsson for valuable discussions. This work used supercomputing resources with allocation award TG-MCB180049 through the Extreme Science and Engineering Discovery Environment (XSEDE), which is supported by National Science Foundation grant number ACI-1548562, and project M2874 through the National Energy Research Scientific Computing Center (NERSC), which is a U.S. Department of Energy Office of Science User Facility operated under Contract No. DE-AC02-05CH11231, and the Research Computing Cluster at the University of Kansas. This work was supported by the American Heart Association (Award 17SDG33370094), the National Institutes of Health (R01GM132572) and the startup funding in the College of Liberal Arts and Sciences at the University of Kansas.

## References

[1] A.L. Hopkins, C.R. Groom, The druggable genome, Nat. Rev. Drug Discovery, 1 (2002) 727.

[2] R. Lappano, M. Maggiolini, G protein-coupled receptors: novel targets for drug discovery in cancer, Nat. Rev. Drug Discovery, 10 (2011) 47.

[3] K.A. Jacobson, New paradigms in GPCR drug discovery, Biochem. Pharmacol., 98 (2015) 541–555.

[4] K.A. Jacobson, Z.-G. Gao, Adenosine receptors as therapeutic targets, Nat. Rev. Drug Discovery, 5 (2006) 247.

[5] R.F. Bruns, J.H. Fergus, Allosteric enhancement of adenosine A1 receptor binding and function by 2-amino-3-benzoylthiophenes, Mol. Pharmacol., 38 (1990) 939–949.

[6] R.F. Bruns, J.H. Fergus, L.L. Coughenour, G.G. Courtland, T.A. Pugsley, J.H. Dodd, F.J. Tinney, Structure-activity relationships for enhancement of adenosine A1 receptor binding by 2-amino-3-benzoylthiophenes, Mol. Pharmacol., 38 (1990) 950–958.

[7] R. Romagnoli, P.G. Baraldi, A.P. Ijzerman, A. Massink, O. Cruz-Lopez, L.C. Lopez-Cara, G. Saponaro, D. Preti, M. Aghazadeh Tabrizi, S. Baraldi, A.R. Moorman, F. Vincenzi, P.A. Borea, K. Varani, Synthesis and Biological Evaluation of Novel Allosteric Enhancers of the A1 Adenosine Receptor Based on 2-Amino-3-(4′-Chlorobenzoyl)-4-Substituted-5-Arylethynyl Thiophene, J. Med. Chem., 57 (2014) 7673–7686.

[8] P.G. Baraldi, A.N. Zaid, I. Lampronti, F. Fruttarolo, M.G. Pavani, M.A. Tabrizi, J.C. Shryock, E. Leung, R. Romagnoli, Synthesis and biological effects of a new series of 2-amino-3-benzoylthiophenes as allosteric enhancers of A1-adenosine receptor, Bioorg. Med. Chem. Lett., 10 (2000) 1953–1957.

[9] C.E. Tranberg, A. Zickgraf, B.N. Giunta, H. Luetjens, H. Figler, L.J. Murphree, R. Falke, H. Fleischer, J. Linden, P.J. Scammells, R.A. Olsson, 2-Amino-3-aroyl-4,5-alkylthiophenes: Agonist Allosteric Enhancers at Human A1 Adenosine Receptors, J. Med. Chem., 45 (2002) 382–389.

[10] L. Aurelio, C. Valant, B.L. Flynn, P.M. Sexton, A. Christopoulos, P.J. Scammells, Allosteric Modulators of the Adenosine A1 Receptor: Synthesis and Pharmacological Evaluation of 4-Substituted 2-Amino-3-benzoylthiophenes, J. Med. Chem., 52 (2009) 4543–4547.

[11] L. Aurelio, C. Valant, H. Figler, B.L. Flynn, J. Linden, P.M. Sexton, A. Christopoulos, P.J. Scammells, 3- and 6-Substituted 2-amino-4,5,6,7-tetrahydrothieno[2,3-c]pyridines as A1 adenosine receptor allosteric modulators and antagonists, Bioorg. Med. Chem., 17 (2009) 7353–7361.

[12] L. Aurelio, A. Christopoulos, B.L. Flynn, P.J. Scammells, P.M. Sexton, C. Valant, The synthesis and biological evaluation of 2-amino-4,5,6,7,8,9-hexahydrocycloocta[b]thiophenes as allosteric modulators of the A1 adenosine receptor, Bioorg. Med. Chem. Lett., 21 (2011) 3704–3707.

[13] C. Valant, L. Aurelio, S.M. Devine, T.D. Ashton, J.M. White, P.M. Sexton, A. Christopoulos, P.J. Scammells, Synthesis and Characterization of Novel 2-Amino-3-benzoylthiophene Derivatives as Biased Allosteric Agonists and Modulators of the Adenosine A1 Receptor, J. Med. Chem., 55 (2012) 2367–2375.

[14] S.J. Hill, L.T. May, B. Kellam, J. Woolard, Allosteric interactions at adenosine A1 and A3 receptors: new insights into the role of small molecules and receptor dimerization, Br. J. Pharmacol., 171 (2014) 1102–1113.

[15] H. Lütjens, A. Zickgraf, H. Figler, J. Linden, R.A. Olsson, P.J. Scammells, 2-Amino-3-benzoylthiophene Allosteric Enhancers of A1 Adenosine Agonist Binding: New 3, 4-, and 5-Modifications, J. Med. Chem., 46 (2003) 1870–1877.

[16] X. Li, D. Conklin, H.-L. Pan, J.C. Eisenach, Allosteric Adenosine Receptor Modulation Reduces Hypersensitivity Following Peripheral Inflammation by a Central Mechanism, J. Pharmacol. Exp. Ther., 305 (2003) 950–955.

[17] S.R. Childers, X. Li, R. Xiao, J.C. Eisenach, Allosteric modulation of adenosine A1 receptor coupling to G-proteins in brain, J. Neurochem., 93 (2005) 715–723.

[18] J. Rahuel, V. Rasetti, J. Maibaum, H. Rüeger, R. Göschke, N. Cohen, S. Stutz, F. Cumin, W. Fuhrer, J. Wood, Structure-based drug design: the discovery of novel nonpeptide orally active inhibitors of human renin, Chem. Biol., 7 (2000) 493–504.

[19] J. Wang, A. Bhattarai, W.I. Ahmad, T.S. Farnan, K.P. John, Y. Miao, Chapter 15 - Computer-aided GPCR drug discovery, in: B. Jastrzebska, P.S.H. Park (Eds.) GPCRs, Academic Press, 2020, pp. 283–293.

[20] S.W. Kaldor, V.J. Kalish, J.F. Davies, B.V. Shetty, J.E. Fritz, K. Appelt, J.A. Burgess, K.M. Campanale, N.Y. Chirgadze, D.K. Clawson, B.A. Dressman, S.D. Hatch, D.A. Khalil, M.B. Kosa, P.P. Lubbehusen, M.A. Muesing, A.K. Patick, S.H. Reich, K.S. Su, J.H. Tatlock, Viracept (Nelfinavir Mesylate, AG1343): A Potent, Orally Bioavailable Inhibitor of HIV-1 Protease, J. Med. Chem., 40 (1997) 3979–3985.

[21] H.A. Carlson, K.M. Masukawa, J.A. McCammon, Method for Including the Dynamic Fluctuations of a Protein in Computer-Aided Drug Design, J. Phys. Chem. A, 103 (1999) 10213–10219.

[22] W. Evangelista Falcon, S.R. Ellingson, J.C. Smith, J. Baudry, Ensemble Docking in Drug Discovery: How Many Protein Configurations from Molecular Dynamics Simulations are Needed To Reproduce Known Ligand Binding?, J. Phys. Chem. B, 123 (2019) 5189–5195.

[23] R.E. Amaro, J. Baudry, J. Chodera, Ö. Demir, J.A. McCammon, Y. Miao, J.C. Smith, Ensemble Docking in Drug Discovery, Biophys. J., 114 (2018) 2271–2278.

[24] J.-H. Lin, A.L. Perryman, J.R. Schames, J.A. McCammon, Computational Drug Design Accommodating Receptor Flexibility: The Relaxed Complex Scheme, J. Am. Chem. Soc., 124 (2002) 5632–5633.

[25] J.R. Schames, R.H. Henchman, J.S. Siegel, C.A. Sotriffer, H. Ni, J.A. McCammon, Discovery of a novel binding trench in HIV integrase, J. Med. Chem., 47 (2004) 1879–1881.

[26] R.E. Amaro, A. Schnaufer, H. Interthal, W. Hol, K.D. Stuart, J.A. McCammon, Discovery of drug-like inhibitors of an essential RNA-editing ligase in Trypanosoma brucei, Proc. Natl. Acad. Sci. USA, 105 (2008) 17278–17283.

[27] K.M. Haynes, N. Abdali, V. Jhawar, H.I. Zgurskaya, J.M. Parks, A.T. Green, J. Baudry, V.V. Rybenkov, J.C. Smith, J.K. Walker, Identification and Structure–Activity Relationships of Novel Compounds that Potentiate the Activities of Antibiotics in Escherichia coli, J. Med. Chem., 60 (2017) 6205–6219.

[28] K. Kapoor, N. McGill, C.B. Peterson, H.V. Meyers, M.N. Blackburn, J. Baudry, Discovery of Novel Nonactive Site Inhibitors of the Prothrombinase Enzyme Complex, J. Chem. Inf. Model., 56 (2016) 535–547.

[29] H.A. Velazquez, D. Riccardi, Z. Xiao, L.D. Quarles, C.R. Yates, J. Baudry, J.C. Smith, Ensemble docking to difficult targets in early-stage drug discovery: Methodology and application to fibroblast growth factor 23, Chem. Biol. Drug Des., 91 (2018) 491–504.

[30] W. Evangelista, R.L. Weir, S.R. Ellingson, J.B. Harris, K. Kapoor, J.C. Smith, J. Baudry, Ensemble-based docking: From hit discovery to metabolism and toxicity predictions, Bioorg. Med. Chem., 24 (2016) 4928–4935.

[31] M. Pi, K. Kapoor, R. Ye, J.C. Smith, J. Baudry, L.D. Quarles, GPCR6A Is a Molecular Target for the Natural Products Gallate and EGCG in Green Tea, Mol. Nutr. Food Res., 62 (2018) 1700770.

[32] H. He, R.L. Weir, J.J. Toutounchian, J. Pagadala, J.J. Steinle, J. Baudry, D.D. Miller, C.R. Yates, The quinic acid derivative KZ-41 prevents glucose-induced caspase-3 activation in retinal endothelial cells through an IGF-1 receptor dependent mechanism, PLOS ONE, 12 (2017) e0180808.

[33] X.-P. Huang, J. Karpiak, W.K. Kroeze, H. Zhu, X. Chen, S.S. Moy, K.A. Saddoris, V.D. Nikolova, M.S. Farrell, S. Wang, T.J. Mangano, D.A. Deshpande, A. Jiang, R.B. Penn, J. Jin, B.H. Koller, T. Kenakin, B.K. Shoichet, B.L. Roth, Allosteric ligands for the pharmacologically dark receptors GPR68 and GPR65, Nature, 527 (2015) 477–483.

[34] Y. Miao, D.A. Goldfeld, E.V. Moo, P.M. Sexton, A. Christopoulos, J.A. McCammon, C. Valant, Accelerated structure-based design of chemically diverse allosteric modulators of a muscarinic G protein-coupled receptor, Proc. Natl. Acad. Sci. USA, 113 (2016) E5675–E5684.

[35] A. Glukhova, D.M. Thal, A.T. Nguyen, E.A. Vecchio, M. Jörg, P.J. Scammells, L.T. May, P.M. Sexton, A. Christopoulos, Structure of the adenosine A1 receptor reveals the basis for subtype selectivity, Cell, 168 (2017) 867–877. e813.

[36] C.J. Draper-Joyce, M. Khoshouei, D.M. Thal, Y.-L. Liang, A.T. Nguyen, S.G. Furness, H. Venugopal, J.-A. Baltos, J.M. Plitzko, R. Danev, Structure of the adenosine-bound human adenosine A 1 receptor–G i complex, Nature, 558 (2018) 559.

[37] R. Romagnoli, P.G. Baraldi, M.A. Tabrizi, S. Gessi, P.A. Borea, S. Merighi, Allosteric Enhancers of A1 Adenosine Receptors: State of the Art and New Horizons for Drug Development, Curr. Med. Chem., 17 (2010) 3488–3502.

[38] A.T.N. Nguyen, E.A. Vecchio, T. Thomas, T.D. Nguyen, L. Aurelio, P.J. Scammells, P.J. White, P.M. Sexton, K.J. Gregory, L.T. May, A. Christopoulos, Role of the Second Extracellular Loop of the Adenosine A1 Receptor on Allosteric Modulator Binding, Signaling, and Cooperativity, Mol. Pharmacol., 90 (2016) 715–725.

[39] A.T.N. Nguyen, J.-A. Baltos, T. Thomas, T.D. Nguyen, L.L. Muñoz, K.J. Gregory, P.J. White, P.M. Sexton, A. Christopoulos, L.T. May, Extracellular Loop 2 of the Adenosine A1 Receptor Has a Key Role in Orthosteric Ligand Affinity and Agonist Efficacy, Mol. Pharmacol., 90 (2016) 703–714.

[40] Y. Miao, A. Bhattarai, A.T.N. Nguyen, A. Christopoulos, L.T. May, Structural Basis for Binding of Allosteric Drug Leads in the Adenosine A1 Receptor, Sci. Rep., 8 (2018) 16836.

[41] A. Bhattarai, J. Wang, Y. Miao, G-Protein-Coupled Receptor–Membrane Interactions Depend on the Receptor Activation State, J. Comput. Chem., 0 (2019).

[42] J. Wang, Y. Miao, Mechanistic Insights into Specific G Protein Interactions with Adenosine Receptors, J. Phys. Chem. B, 123 (2019) 6462–6473.

[43] F. Vincenzi, M. Targa, R. Romagnoli, S. Merighi, S. Gessi, P.G. Baraldi, P.A. Borea, K. Varani, TRR469, a potent A1 adenosine receptor allosteric modulator, exhibits anti-nociceptive properties in acute and neuropathic pain models in mice, Neuropharmacology, 81 (2014) 6–14.

[44] M. Kimatrai-Salvador, P.G. Baraldi, R. Romagnoli, Allosteric modulation of A1-adenosine receptor: a review, Drug Discovery Today: Technol., 10 (2013) e285–e296.

[45] K.A. Jacobson, Z.-G. Gao, A. Göblyös, A.P. Ijzerman, Chapter 7 - Allosteric Modulation of Purine and Pyrimidine Receptors, in: K.A. Jacobson, J. Linden (Eds.) Adv. Pharmacol., vol. 61, Academic Press, 2011, pp. 187–220.

[46] H. Figler, R.A. Olsson, J. Linden, Allosteric Enhancers of A<sub>1</sub> Adenosine Receptors Increase Receptor-G Protein Coupling and Counteract Guanine Nucleotide Effects on Agonist Binding, Mol. Pharmacol., 64 (2003) 1557–1564.

[47] M.M. Mysinger, M. Carchia, J.J. Irwin, B.K. Shoichet, Directory of Useful Decoys, Enhanced (DUD-E): Better Ligands and Decoys for Better Benchmarking, J. Med. Chem., 55 (2012) 6582–6594.

[48] G.M. Morris, R. Huey, W. Lindstrom, M.F. Sanner, R.K. Belew, D.S. Goodsell, A.J. Olson, AutoDock4 and AutoDockTools4: Automated docking with selective receptor flexibility, J. Comput. Chem., 30 (2009) 2785–2791.

[49] S. Forli, R. Huey, M.E. Pique, M.F. Sanner, D.S. Goodsell, A.J. Olson, Computational protein–ligand docking and virtual drug screening with the AutoDock suite, Nat. Protoc., 11 (2016) 905.

[50] P.A. Ravindranath, S. Forli, D.S. Goodsell, A.J. Olson, M.F. Sanner, AutoDockFR: advances in protein-ligand docking with explicitly specified binding site flexibility, PLoS Comput. Biol., 11 (2015) e1004586.

[51] D.A. Pearlman, P.S. Charifson, Improved scoring of ligand− protein interactions using OWFEG free energy grids, J. Med. Chem., 44 (2001) 502–511.

[52] Y. Miao, V.A. Feher, J.A. McCammon, Gaussian Accelerated Molecular Dynamics: Unconstrained Enhanced Sampling and Free Energy Calculation, J. Chem. Theory Comput., 11 (2015) 3584–3595.

[53] Y. Miao, J.A. McCammon, Gaussian Accelerated Molecular Dynamics: Theory, Implementation, and Applications, Annu. Rep. Comput. Chem., 13 (2017) 231–278.

[54] Y. Miao, Acceleration of biomolecular kinetics in Gaussian accelerated molecular dynamics, J. Chem. Phys., 149 (2018) 072308.

[55] D. Hamelberg, J. Mongan, J.A. McCammon, Accelerated molecular dynamics: a promising and efficient simulation method for biomolecules, J. Chem. Phys., 120 (2004) 11919–11929.

[56] T. Shen, D. Hamelberg, A statistical analysis of the precision of reweighting-based simulations, J. Chem. Phys., 129 (2008) 034103.

[57] D.S.C. D.A. Case, T.E. Cheatham, III, T.A. Darden, R.E. Duke, T.J. Giese, H. Gohlke, A.W. Goetz, D. Greene, N. Homeyer, S. Izadi, A. Kovalenko, T.S. Lee, S. LeGrand, P. Li, C. Lin, J. Liu, T. Luchko, R. Luo, D. Mermelstein, K.M. Merz, G. Monard, H. Nguyen, I. Omelyan, A. Onufriev, F. Pan, R. Qi, D.R. Roe, A. Roitberg, C. Sagui, C.L. Simmerling, W.M. Botello-Smith, J. Swails, R.C. Walker, J. Wang, R.M. Wolf, X. Wu, L. Xiao, D.M. York and P.A. Kollman (2018), AMBER 2018, University of California, San Francisco.

[58] Y.T. Pang, Y. Miao, Y. Wang, J.A. McCammon, Gaussian Accelerated Molecular Dynamics in NAMD, J. Chem. Theory Comput., 13 (2017) 9–19.

[59] Y. Miao, W. Sinko, L. Pierce, D. Bucher, R.C. Walker, J.A. McCammon, Improved Reweighting of Accelerated Molecular Dynamics Simulations for Free Energy Calculation, J. Chem. Theory Comput., 10 (2014) 2677–2689.

[60] D. Hamelberg, C.A. de Oliveira, J.A. McCammon, Sampling of slow diffusive conformational transitions with accelerated molecular dynamics, J. Chem. Phys., 127 (2007) 155102.

[61] B. Roux, The calculation of the potential of mean force using computer simulations, Comput. Phys. Commun., 91 (1995) 275–282.

[62] W. Humphrey, A. Dalke, K. Schulten, VMD: visual molecular dynamics, J. Mol. Graph., 14 (1996) 33–38, 27–38.

[63] K. Vanommeslaeghe, A.D. MacKerell, Jr., CHARMM additive and polarizable force fields for biophysics and computer-aided drug design, Biochim. Biophys. Acta., 1850 (2015) 861–871.

[64] K. Vanommeslaeghe, E. Hatcher, C. Acharya, S. Kundu, S. Zhong, J. Shim, E. Darian, O. Guvench, P. Lopes, I. Vorobyov, A.D. Mackerell, Jr., CHARMM general force field: A force field for drug-like molecules compatible with the CHARMM all-atom additive biological force fields, J. Comput. Chem., 31 (2010) 671–690.

[65] K. Kappel, Y. Miao, J.A. McCammon, Accelerated molecular dynamics simulations of ligand binding to a muscarinic G-protein-coupled receptor, Q. Rev. Biophys., 48 (2015) 479–487.

[66] J.C. Phillips, R. Braun, W. Wang, J. Gumbart, E. Tajkhorshid, E. Villa, C. Chipot, R.D. Skeel, L. Kale, K. Schulten, Scalable molecular dynamics with NAMD, J. Comput. Chem., 26 (2005) 1781–1802.

[67] D.R. Roe, T.E. Cheatham, 3rd, PTRAJ and CPPTRAJ: Software for Processing and Analysis of Molecular Dynamics Trajectory Data, J. Chem. Theory Comput., 9 (2013) 3084–3095.

[68] G.M. Morris, D.S. Goodsell, R.S. Halliday, R. Huey, W.E. Hart, R.K. Belew, A.J. Olson, Automated docking using a Lamarckian genetic algorithm and an empirical binding free energy function, J. Comput. Chem., 19 (1998) 1639–1662.

[69] C.F. Wong, Flexible receptor docking for drug discovery, Expert Opin. Drug Discov., 10 (2015) 1189–1200.

